# Inhibition of Osteoblast Proliferation and Migration by Exogenous and Endogenous Formaldehyde

**DOI:** 10.1101/584052

**Authors:** Xu Teng, Wei Huang, Hefeng Yu, Pei Wang, Weishi Li, Zhongqiang Chen, Dongwei Fan

## Abstract

Exogenous and endogenous formaldehyde (FA) plays an important role in cell growth and migration; however, its potential role in osteoblasts remains largely unclear. Cell counting kit-8 (CCK-8) and wound healing assays revealed that FA exposure at naturally occurring concentrations inhibited the proliferation and migration of mouse preosteoblast MC3T3-E1 cells. Moreover, RNA sequencing (RNA-seq) analysis revealed that FoxO1 signaling pathway components displayed distinct expression patterns upon FA exposure, reflected through significant enrichment of cell migration. In particular, FoxO1 Sirt1 and FA-induced related protein expression which were closely with cell proliferation and migration were confirmed by western blotting. The present results indicate that the FoxO1 pathway is involved in FA-induced inhibition of cell growth and migration.

## 1. Introduction

Osteoblast growth and migration are essential for not only bone metabolism, including bone remodeling and responses to mechanical loading, but also bone pathological processes including bone repair after fractures and during osteoporosis [1–3].

Formaldehyde (FA) is one of the primary factors causing sick building syndrome. FA enters the body through inhalation from building materials, furniture, tobacco smoke, e-cigarettes, the sweetener aspartame, and, most directly, accidental consumption of methanol and insulation materials [4,5]. The WHO-recommended limit for indoor FA is 0.1 mg/m^3^. However, indoor FA concentrations often exceed this recommended threshold. The average concentration of occupational FA exposure in Chinese factories is 1.37 mg/m^3^, which also exceeds the recommended threshold [6]. Simultaneously, FA is present in the natural environment, thus making it difficult to prevent FA exposure [7].

Furthermore, FA is produced endogenously, being found ubiquitously in cells owing to enzymatic oxidative demethylation by-products. Histones of the KDM1/JMJC or ABH enzyme family and the RNA and DNA demethylation machinery produce FA in the nucleus [8,9]. Moreover, FA can be endogenously produced by the action of neutrophil enzyme myeloperoxidase and N-demethylation, a common biochemical phenomenon. Endogenous FA is present in the blood at 50-100 μM [10–12].

Therefore, because of its abundance and chemical properties, formaldehyde can pose a significant risk to genomic integrity. However, limited information is available regarding the mechanisms underlying protection against FA at the cellular and organismal levels. It is unclear whether FA affects bone cells, especially osteoblasts and their proliferation and migration, which play essential roles in bone-associated diseases, e.g., bone repair during fracture and osteoporosis.

This study aimed to characterize the effect of formaldehyde on osteoblast growth and migration. First, we investigated the effect of FA on mouse preosteoblast MC3T3-E1 cells; thereafter, we attempted to determine the key signaling pathway affected in osteoblasts upon FA exposure via genome-wide transcriptional profiling analysis using RNA-seq.

## 2. Materials and methods

### 2.1. Cell culture

MC3T3-E1 cells were purchased from the ATCC (Manassas, VA, USA). Cells were cultured in a humidified atmosphere of 5% CO_2_ and 95% air at 37°C in an alpha modification of Eagle’s medium (Invitrogen Life Technologies, Carlsbad, CA, USA) supplemented with 10% FBS, 100 U/mL penicillin, and 0.1 mg/mL streptomycin[13].

### 2.2. Cell proliferation

Cell proliferation was assessed via a colorimetric tetrazolium salt-based assay, i.e., the cell counting-8 (CCK-8) assay. MC3T3-E1 cells were seeded at a density of 5000 cells/well into a 96-well plate and cultured in the absence or presence of FA at different concentrations (100μM - 500 μM). Thereafter, the CCK-8 solution (Dojindo) was added to each well, and the plate was incubated at 37°C in a CO_2_ incubator for 3h. The absorbance was then determined at 450 nm, using a microplate reader (Bio-Rad, Hercules, CA, USA).

### 2.3. Wound-healing assay

Cells were seeded in a 6-well plate the day before the assay, and cells were pre-treated with FA for 2 h, 6 h or 12 h before the wounds were inflicted on the cell monolayer. The wound was generated by scratching the cell monolayer with a pipette tip, and images were acquired at 0, 12, and 24 h after wounding. The degree of cell migration required to close the wound was analyzed using ImageJ software (National Institutes of Health, Bethesda, MD, USA).

### 2.4. RNA preparation

Total RNA was extracted using TRIzol reagent (Invitrogen, CA, USA, 15596026) on dry ice and processed in accordance with the manufacturer’s instructions. To eliminate DNA, an aliquot of total RNA was treated with RQ1 RNase-Free DNase (Promega, WI, USA, M6101), followed by phenol/chloroform/isoamyl alcohol extraction and chloroform/isoamyl alcohol extraction, using Phase Lock Gel Light tubes (5 PRIME 2302800), followed by ethanol precipitation. Precipitated RNA was stored at −20°C until use.

### 2.5. Illumina RNA-Seq library construction

Total RNA (20 mg) was used for poly(A)t selection using oligo(dT) magnetic beads (Invitrogen, CA, USA, 610-02), eluted in water, and used to generate an RNA-seq library, using the ScriptSeq kit (Epicentre, CA, USA SS10906). Libraries were amplified via polymerase chain reaction for 12–15 cycles and sequenced in two lanes on the HiSeq 2000 platform at BGI Genome Center (Shenzhen, China).

### 2.6. RNA sequencing

High-throughput sequencing was performed as paired-end 100 sequencing, using a HiSeq 2000 system (Illumina, San Diego, CA, USA). RNA-Seq reads were mapped using TopHat software (http://tophat.cbcb.umd.edu) to generate the alignment file, which was then used to assemble transcripts, estimate abundance, and detect differential expression of genes/isoforms, using cufflinks. Gene classification was based on searches in the DAVID (david.abcc.ncifcrf.gov) and MEDLINE (www.ncbi.nlm.nih.gov) databases.

### 2.7. Illumina data analysis

Raw reads were aligned to the B73 reference genome (RefGen v3), using TopHat 2.0.8 and STAR, with the minimum intron length set at 20 bp and the maximum intron length set at 50 kb, with default settings for other parameters. Genes and isoforms were quantified using cufflinks version 2.2.1, using the GTF annotation file generated by PacBio sequencing. To reduce transcriptional noise, only isoforms/genes with FPKM values of ≥0.01 (based on gene coverage saturation analysis) were included.

### 2.8. In silico analysis

To identify the pathways of intersecting genes, pathway enrichment analysis was performed using the Kyoto Encyclopedia of Genes and Genomes (KEGG, http://www.genome.ad.jp/kegg/). This analysis provides a better understanding of gene expression as a complete network. The Fisher’s exact test and Chi-square test were performed, and the threshold for statistical significance was defined by a P-value of <0.05 and the false discovery rate. Furthermore, we performed gene-set enrichment analysis, using GSEA v2.2.2 software and a c5.all. v5.1. symbols.gmt (gene ontology)-gene set matrix.

### 2.9. Western blot analysis

MC3T3-E1 cells were cultured in a 10-cm culture dish at 5 × 10^5^ cells/mL at 37°C in 5% CO_2_ in medium supplemented with FA at different concentrations. Thereafter, cells were harvested, and extracts were cleared via centrifugation (Thermo Scientific, CA, USA). Protein concentrations were determined using the BCA protein assay kit (Thermo Scientific, CA, USA). Protein samples (50 μg each) were separated via SDS-PAGE (12% resolving gel) and then transferred to polyvinylidene difluoride membranes. The membranes were blocked with blocking solution containing 3% BSA and 0.1% Tween-20 for 30 min and probed overnight at 4°C with the following primary antibodies: anti-FoxO1, anti-SIRT1, anti-PRMT6, anti-SUZ12, anti-EZH2 (1:500 each) (Merck Millipore, Darmstadt, Germany), and anti-GAPDH (1:5000) (Bioworld Technology, Saint Louis Park, MN, USA). The membrane was then probed with HRP-labeled goat anti-rabbit IgG (H + L) secondary antibody (1:1000; KPL, Milford, MA, USA) for 1 h. Finally, protein bands were detected using DAB reagent. Relative protein expression levels were determined using Quantity One (Bio-Rad).

### 2.10. Statistical analysis

Unless indicated otherwise, data are presented as mean values ± standard deviation (SD) values and analyzed using one-way ANOVA, followed by the Dunnett multiple comparison test for post hoc analysis. A *p* value <0.05 was considered statistically significant. Data were analyzed using SPSS11 (Chicago, IL, USA) statistical software.

## 3. Results

### 3.1. Effect of FA on osteoblast proliferation

The effect of FA on proliferation in MC3T3-E1 osteoblasts was determined using the CCK-8 assay. As shown in Fig. 1, cell proliferation was observed upon treatment with FA at different concentrations for 2 h (Fig. 1. upper panel) and 6 h (Fig. 1. lower panel). Typically, each group treated with 100 μM, 150 μM, 200 μM, 300 μM, and 500 μM FA markedly inhibited proliferation in comparison with untreated cells, and longer the duration of FA treatment, stronger the inhibition of cell proliferation. These results notably indicate the strong inhibitory effect of FA on MC3T3-E1 cells through reduced cell viability in a dose-dependent manner within 0–96 h (Fig. 1).

**Fig. 1.**
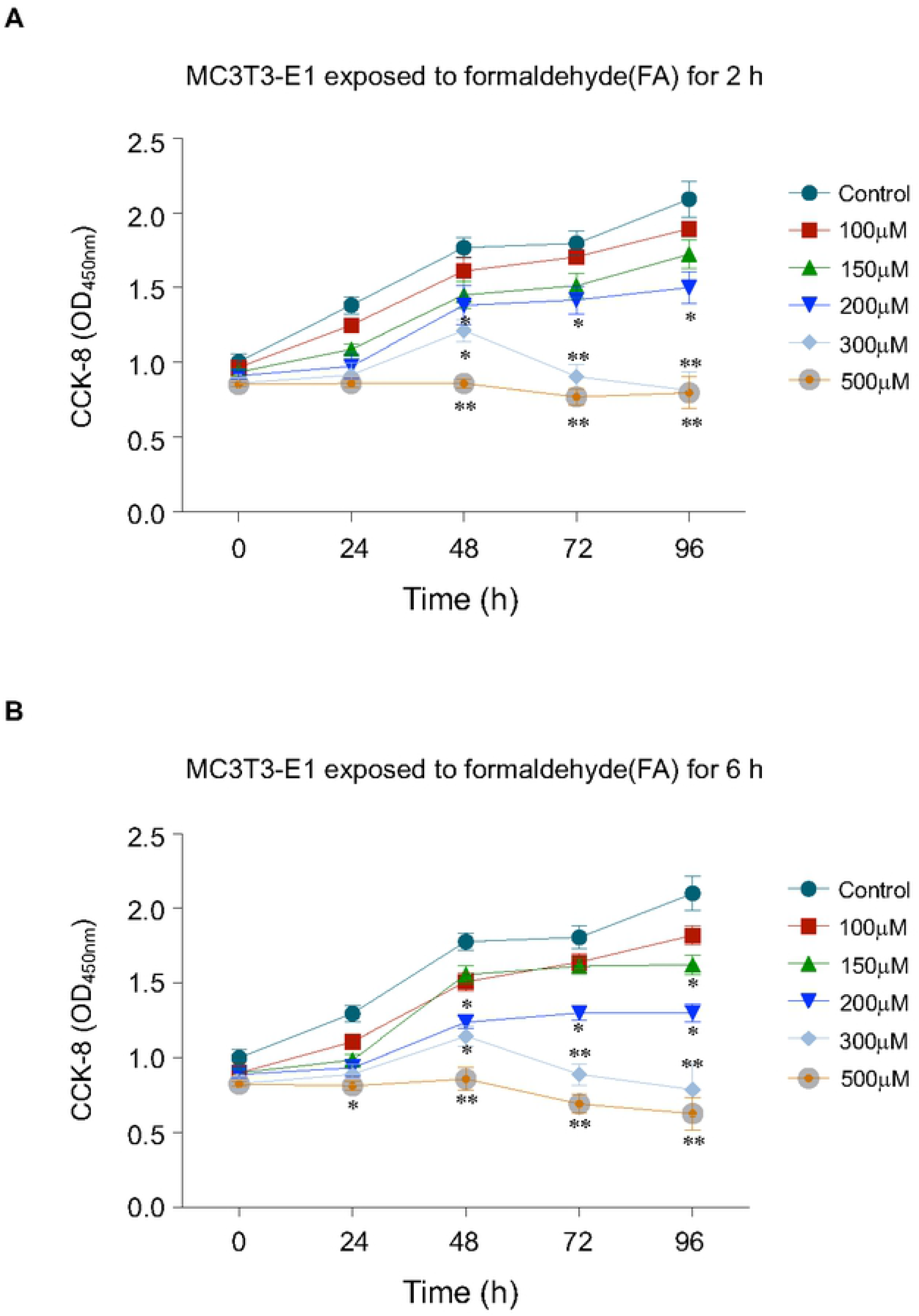
Effects of formaldehyde (FA) on cell viability in MC3T3-E1. MC3T3-E1 cells were treated for 2 h (A) and 6 h (B) with the control or with FA at different concentrations, and cell viability was examined using the cell counting kit-8 assay at 0, 24, 48, 72, and 96 h. The results are expressed as the mean ± SD values of at least three independent experiments. \**p* < 0.05, \*\**p* < 0.01.

### 3.2. Effect of FA on osteoblast migration

We investigated the effect of FA on the migration of MC3T3-E1 cells, using a wound-healing assay. Upon treatment of cells for 2 h, 6 h and 12 h with 100 μM, 150 μM, 200 μM, 300 μM, and 500 μM FA, respectively, wounds were inflicted on the surface of the confluent cellular monolayer surface, and cells were allowed to migrate for 12 and 24 h. The wound healing assays showed that FA significantly decreased the rate of wound closure of MC3T3-E1 cells in comparison with the control at 12 h and 24 h of treatment with 200 μM to 500 μM FA (*p* < 0.05, Dunnett comparison test) (Fig. 2). In addition, whether the cells were treated for 2 h, 6 h, or 12 h, the wound closure rate of MC3T3-E1 cells decreased with the increase of FA dose, showing a dose-dependent inhibition of cell migration.

**Fig. 2.**
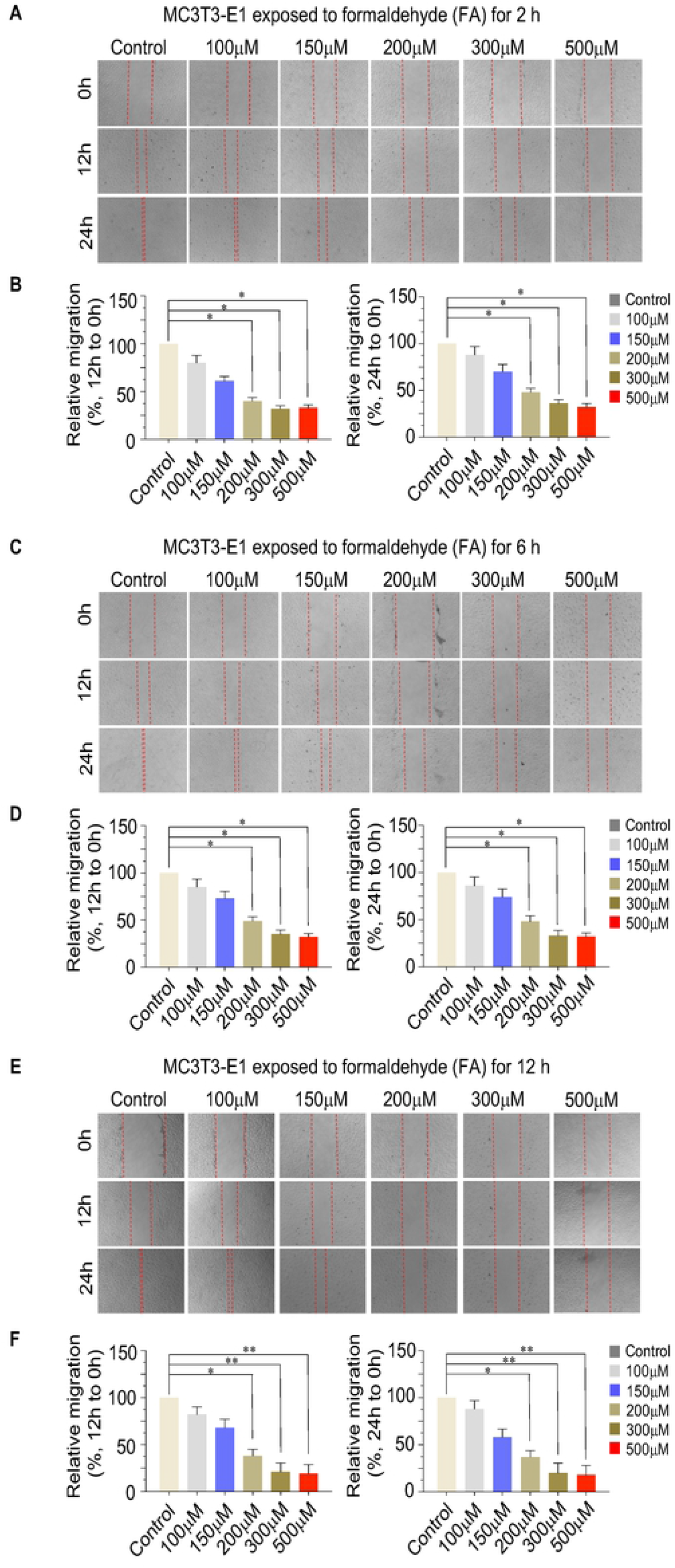
Effects of formaldehyde (FA) on cell migration in MC3T3-E1. Cells were treated for 2 h (A), 6 h (C) and 12 h (E) with the control or with FA at different concentrations, and cell migration was examined via a wound-healing assay. Statistical analysis of migrations in each group with 12 h (left panel) and 24 h (right panel) were shown in (B), (D) and (F) panel. The results are expressed as mean ± SD values of at least three independent experiments. \**p* < 0.05, \*\**p* < 0.01.

### 3.3. Identification of the candidate signaling pathway involved in cell proliferation and migration after FA treatment

To explore the potential mechanism underlying osteoblast proliferation and migration after FA treatment at the whole-genome level, we performed a genome-wide transcriptional analysis via RNA-seq. For cells treated with each FA concentration, whole-genome transcriptomes were determined. In brief, after treatment of MC3T3-E1 cells with 100 μM and 300 μM FA for 6 h, cells were harvested for RNA extraction and subjected to RNA-seq. Preliminary results indicate that 100 μM and 300 μM FA differentially up- and downregulated numerous genes (Fig. 3A), and the differentially expressed genes were enriched upon KEGG pathway analysis, indicating that these genes are involved in cell adhesion. Signaling pathways such as those involving FoxO, Wnt, autophagy-related proteins, MAPK, mTOR, Hippo, and p53 (Fig. 3B) are involved in cell growth, proliferation, migration, and invasion. The 972 differentially upregulated genes and the 3968 differentially downregulated genes were separately enriched in the KEGG pathway. These upregulated genes were found to be primarily involved in DNA damage repair, cell metabolism, and other signaling pathways, while the downregulated genes were primarily involved in p53, Hippo, autophagy, FoxO, Wnt, TGF-beta, and other signaling pathways (Fig. 3C). Furthermore, analysis of the RNA-seq results via GSEA revealed that the FA-300 group significantly inhibited signaling pathways such as FoxO, p53, Wnt, and mTOR (*p* < 0.05, Dunnett multiple comparison test) (Fig. 3D). These results preliminarily indicate that the *in vitro* MC3T3-E1 osteoblast model of FA treatment is indeed reliable.

**Fig. 3.**
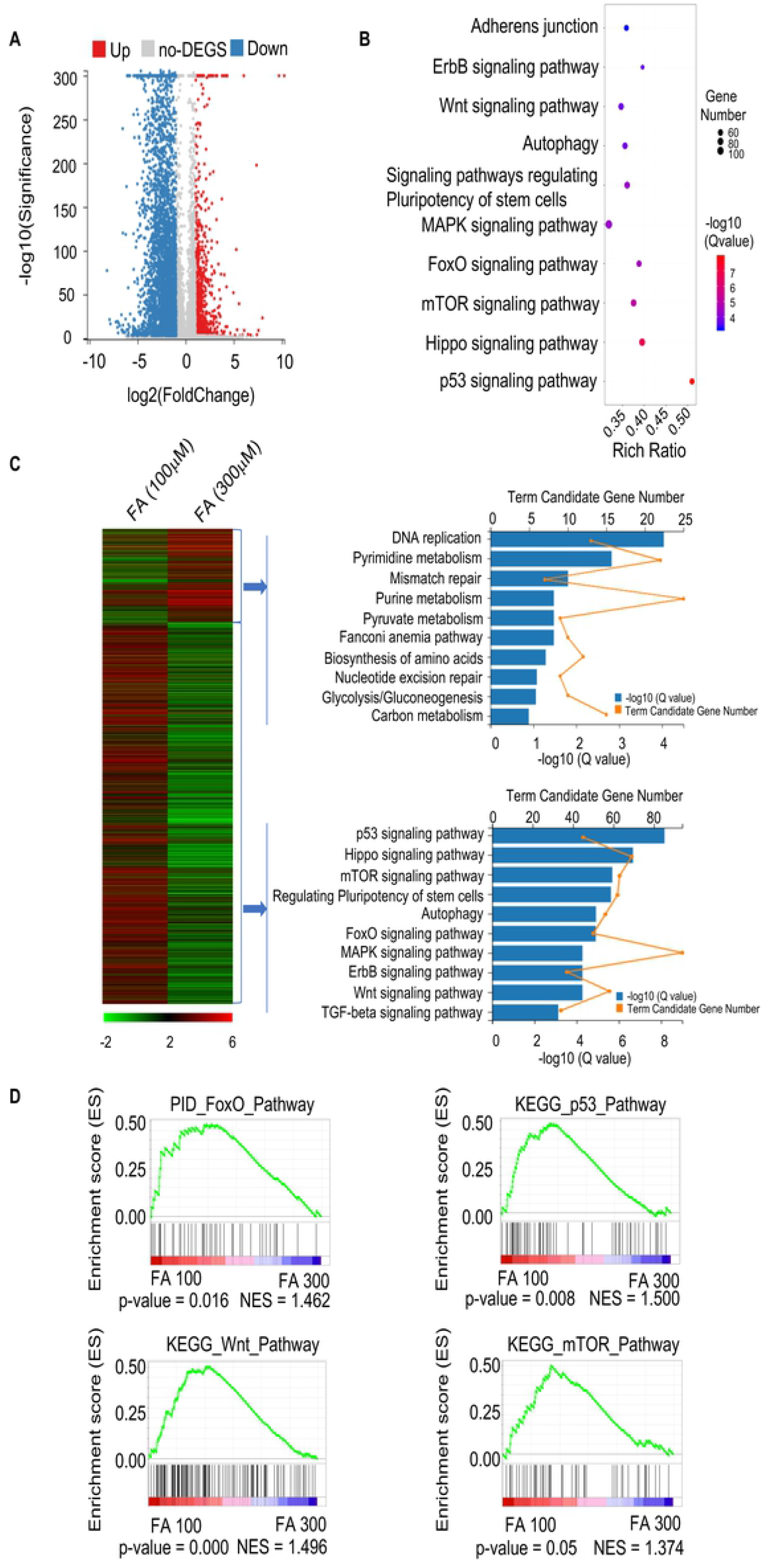
MC3T3-E1 cells treated with 100 μM and 300 μM formaldehyde (FA) for RNA-seq and analysis of RNA-seq results. (A) RNA-seq revealed that 972 genes were upregulated and 3968 genes were downregulated (fold-change > 2) upon treatment with 100 μM and 300 μM FA. (B) Differential gene enrichment in KEGG pathways upon treatment with 100 μM and 300 μM FA, showing that these genes are involved in cell adhesion, Wnt, autophagy, MAPK, FoxO, mTOR, Hippo, and p53 signaling pathways. (C) The 972 differentially upregulated genes and the 3968 differentially downregulated genes were separately enriched in the KEGG pathway. (D) GSEA analysis of RNA-seq results.

### 3.4. Effect of FA on FOXO1/SIRT1 protein expression

To evaluate effect of FA on FOXO1 [14], SIRT1, PRMT6, SUZ12 [15], and EZH2 [16] protein expression levels, which are closely associated with cell proliferation and migration in MC3T3-E1 cells, we treated cells with FA at increasing concentrations from 100 μM to 300 μM. Western blot analysis revealed that FOXO1, SIRT1, PRMT6, SUZ12, and EZH2 were significantly downregulated upon FA treatment (Fig. 4).

**Fig. 4.**
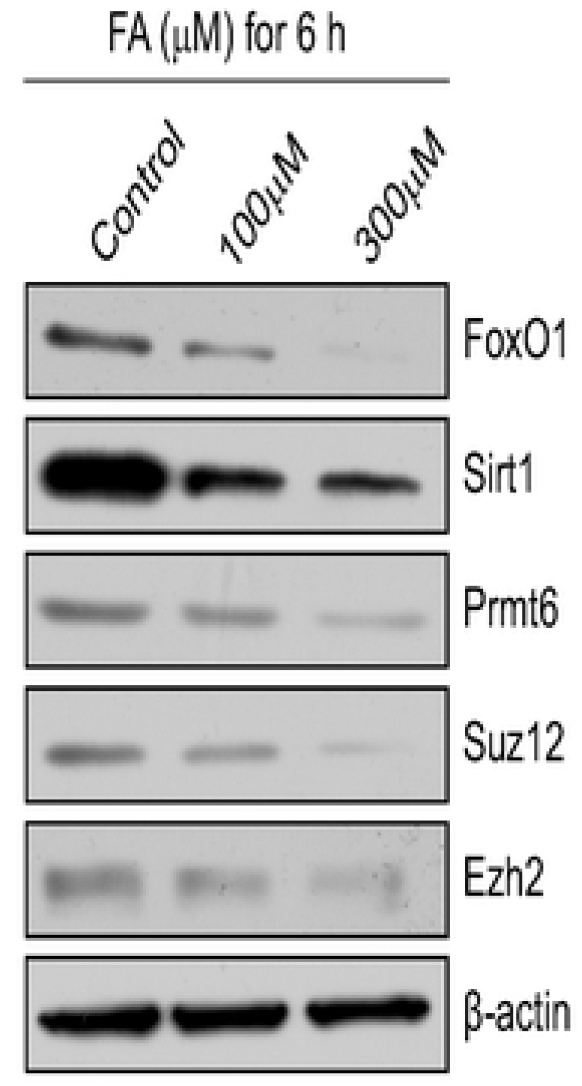
Altered FoxO1 expression levels in MC3T3-E1 cells after FA treatment. MC3T3-E1 were seeded in 6-well plates and treated with formaldehyde (FA). FoxO1 expression levels and FA-induced related protein expression levels were determined via western blot analysis.

## 4. DISCUSSION

To our knowledge, this study is the first to report that FA, a product of cellular metabolism and a ubiquitous environmental toxin, suppressed osteoblastic cell proliferation and decreased cell migration at naturally occurring concentrations. Furthermore, a genome-wide transcriptional analysis via RNA-seq performed to explore the potential mechanism underlying FA stimulation revealed distinct expression patterns of components of the FoxO signaling pathway upon FA treatment, reflected through significantly decreased cell migration and proliferation.

Forkhead box O1 (FoxO1), also known as the forkhead rhabdomyosarcoma transcription factor (FKHR), belongs to the Forkhead box (FOX) family and is a key transcriptional regulator in cell proliferation, differentiation, and migration via a receptor tyrosine kinase signaling pathway [17,18]. Recent studies have reported that osteoblast growth and migration is regulated by FoxO factors [19,20]. FoxO1 is the primary regulator of redox balance and function in osteoblasts among three key members of the FoxO family, i.e., FoxO1, FoxO3a, and FoxO4. Depletion of FoxO1 in osteoblasts reportedly resulted in decreased proliferation and bone volume in FoxO1_Ob_^−/−^ mice [21,22]. In the present study, genome-wide expression profiling revealed that *FOXO1* was downregulated upon FA exposure. Furthermore, using protein semi-quantification techniques, FoxO1 was downregulated at higher FA concentrations.

Transcription-related factors can be integrated into specific DNA sites, and this molecular process is regulated by the configuration changes of chromatin. However, highly condensed chromatin is relieved through chromatin remodeling of itself with the alteration of covalence power between histone and DNA chain, and this mechanism may control gene expression and silencing.

Recently, several studies demonstrated a critical role of the Prmt6, Suz12, Ezh2 that can not only affect cell proliferation and differentiation through chromatin remodeling, but this process is related to FoxO signaling pathway. For example, oxidative stress in cells, activation of SIRT/FOXO/SOD2 signaling pathway, in turn, recruits chromatin remodeling complexes (Prmt6, Suz12, Ezh2) to activate related gene expression[23]. Our genome-wide analysis found that not only the FOXO1 and SIRT1 expression was reduced, but also the expression of chromatin remodeling key proteins including PRMT6, SUZ12, and EZH2 was down-regulated, and further confirmed by Western Blot.

These data indicate that FoxO1 plays an important role in FA-induced inhibition of proliferation and migration in MC3T3-E1 cells, and further studies are intended to clarify its upstream and downstream interacting partners to further clarify the detailed mechanism of action of FA on cell proliferation and migration. In the present study, transcriptomic analysis of cellular responses elucidated the potential underlying mechanism and the dose-dependent response pattern.

Although the present results cannot be extrapolated directly to the *in vivo* setting, further studies are required to identify potentially sensitive target genes, since dysregulated expression of these genes can affect osteogenesis and cause bone disease.

In conclusion, the present results show that FA inhibits cell migration and proliferation in MC3T3-E1 cells. Furthermore, this newly identified effect of FA on cell migration can be harnessed not only to prevent air pollution but also for novel therapeutic applications such as acceleration of wound healing by accumulation of preosteoblast cells at the bone fracture site. Therefore, further studies are required to clarify the effect of FA on FoxO1 and other target molecules mediating FA-induced inhibition of cell migration and proliferation.

## Conflicts of interest

There is no conflict of interests in this work.

## Acknowledgements

*This work was supported by grants from National Natural Science Foundation of China (81672726; 30801156); National Key Research and Development Program of China (2017YFC0113001); Program for Training Capital Science and Technology Leading Talents (Z181100006318003); and Fostering Young Scholars of Peking University Health Science Center (BMU2017PY015)

